# Improving interpretability of deep learning models: splicing codes as a case study

**DOI:** 10.1101/700096

**Authors:** Anupama Jha, Joseph K. Aicher, Deependra Singh, Yoseph Barash

## Abstract

Despite the success and fast adaptation of deep learning models in a wide range of fields, lack of interpretability remains an issue, especially in biomedical domains. A recent promising method to address this limitation is Integrated Gradients (IG), which identifies features associated with a prediction by traversing a linear path from a baseline to a sample. We extend IG with nonlinear paths, embedding in latent space, alternative baselines, and a framework to identify important features which make it suitable for interpretation of deep models for genomics.

## Background

The high accuracy of deep neural networks (DNN) in areas such as computer vision, natural language processing, and robotics has led to the fast adaptation of DNN in biomedical research. In genomics, deep learning models have outperformed previous state-of-the-art methods on tasks such as predicting protein binding sites (Zhou & Troyanskaya, 2015) or mRNA alternative splicing from genomic sequence features (Jha et al., 2017). However, the interpretation of these complex models remains a challenge (Guidotti et al., 2018; Lipton, 2016). Approaches to model interpretation include approximation with simpler models (Ribeiro et al., 2016), identifying the most influential samples (Koh & Liang, 2017), or finding the most relevant features for a specific sample or a task by a variety of metrics (Baehrens et al., 2010). Here we focus on the latter approach, and specifically on the recently-developed Integrated Gradients (IG) (Sundararajan et al., 2017). In this context, interpretability is defined as attributing the prediction of a DNN to its input features. IG, and similarly DeepLIFT (Shrikumar et al., 2017), identify features associated with a model’s prediction with respect to a baseline. The usage of a baseline is attractive as it serves as the model’s proxy to human counterfactual intuition, assigning blame by absence of a feature. IG computes feature attribution by aggregating gradients along a linear path between the sample and the baseline. Compared to other interpretation methods, IG offers two desirable theoretical guarantees motivating its usage. The first is *sensitivity*, which states that for every input and baseline that differ in one feature but have different predictions the method would give a non-zero attribution for that differing feature. The second is *implementation invariance*, which states that regardless of network architecture if two models are functionally equivalent (same output given any input) then their feature attributions should also be equivalent (see more details in (Shrikumar et al., 2017)).

While IG has been shown to excel on the object recognition problems, the approach taken in the original IG work suffers from several limitations. First, IG only provides feature attributions for individual features with respect to a specific sample. There is no mechanism to assess significance of these attributions apart from a non-zero value, and there is no formal mechanism for identifying significant features for a class of interest. Second, they take a linear path between the baseline and the sample. The authors speculate that paths visiting points far-removed from actual points seen in training could lead to attribution artifacts. This concern could also happen for linear paths, particularly in high-dimensional datasets where observed data may lie close to a (hidden) lower-dimensional nonlinear structure in the original feature space (*i.e*., manifold hypothesis). In such a case, a nonlinear path taken close to the observed training data might be preferable. Finally, IG attributions show features that distinguish a sample from an all-zero, or no signal, baseline point. This creates two potential issues for the genomics domain. First, in many genomics applications it is unclear what inputs actually reflect no signal and a zero might not be the appropriate reference point (*e.g*., exon length of 0 is not biologically meaningful). Second, following the counterfactual argument above, features that distinguish two classes of samples are arguably more useful than a no signal baseline because they can help focus on the relevant bits of information. For example, features associated with RNA binding proteins that distinguish differentially included cassette exons in brain from constitutively spliced exons can help shed light on brain-specific RNA splicing regulation.

We address the limitations of IG listed above using RNA alternative splicing (AS) code models as the main usage case (Jha et al., 2017). Briefly, given a triplet of exons in the pre-mRNA, the middle exon (cassette exon) can be either included or skipped in the mature mRNA, giving rise to different isoforms. The model predicts the middle exon’s inclusion levels (Ψ ∈ [0,1]) and differential inclusion (ΔΨ ∈ [−1,1]) between different conditions (e.g. brain vs. liver) as a function of a 1, 357-dimensional feature vector. These features are parsed from genomic regions containing the exon triplet and flanking introns (see Supplementary Tables. S1-S3 and Supplementary Figure. S1 for details about the architecture). The splicing variations (Ψ) used here to train and test the models were derived from RNA-Seq experiments involving six mouse tissues (Hippocampus, Heart, Liver, Lung, Spleen, Thymus) (Keane et al., 2011), and quantified using MAJIQ (Vaquero-Garcia et al., 2016). Specifically, for the results described below, we use a group of cassette exons that are differentially included in hippocampus compared to at least one of the other five tissues in the dataset as the set of AS events of interest and constitutively spliced exons as the baseline class of events.

These splicing code models have several desirable characteristics for analyzing DNN interpretation for genomic tasks. First, the features are an identifiable set representing prior biological knowledge about putative regulatory elements such as known sequence motifs and RNA secondary structure. Moreover, a myriad of models have already been applied to this task, including a mixture of decision trees, bayesian neural networks, naive bayes, and logistic regression (Barash et al., 2010; Xiong et al., 2011). Second, the splicing code model includes embedding in a lower dimension space, a common component in genomic models, allowing us to test the usage of feature embedding for prediction attribution.

In addition to splicing code models, we also use models for the handwritten digit recognition task (LeCun, 1998) to demonstrate the applicability of our ideas in a different, easier to visualize, domain. The digit model predicts the identity of a handwritten digit between 0 and 9 from a 28 × 28 pixel image (see Supplementary Tables. S4-S6 and Supplementary Figure. S2 for details about the architecture).

## Results

### A robust framework for identifying significant features

Training the splicing code models involves the training of an autoencoder or variational autoencoder as the first step. Thus, any downstream analysis should first address the stability of these nonlinear embeddings in a lower dimensional space. Given our interest in paths between points and the advantages demonstrated in other domains for assessing similarities based on local structures (Roweis & Saul, 2000; Wu & Schölkopf, 2007; Wang et al., 2017), we test stability of the embeddings via neighborhood similarity. Specifically, we evaluated several autoencoder and variational autoencoder architectures to conclude the embeddings do not vary much, with an observed spearman rank correlation typically in the range of 0.85 – 0.95 when varying architectures, bootstrapping samples and initializations (See Methods and Supplementary Figure. S3).

Using the latent embedding from the trained autoencoder as input, we train a feed-forward network with similar architecture as (Jha et al., 2017) (see Supplementary Figure. S1 for details about the architecture). The combined encoder and feed-forward network predict the splicing outcome. This combined architecture enables us to run attribution in the original and hidden space. We also address a potential stability issue of the integral approximation performed by IG. Based on this analysis we selected 250 points for the experiments described below (see Supplementary section 3 on path interpolation and Supplementary Figure S4 for details).

To identify significant features for any group of splicing samples, we apply the following procedure. First, we group highly-correlated features (*e.g*., different versions of the FOX splicing factor binding motif) into *meta-features* in order to avoid spreading attributions associated with the same entity (FOX binding in this example). Next, we identify significant features by comparing the relative ranking of a meta-feature’s absolute attribution across samples belonging to a class of interest to those observed for a similarly sized random set of samples using one-sided t-test with Bonferroni correction for multiple testing (an illustration is shown in Fig. 1c). We validate that the resulting empirical distribution of *p*-values is well calibrated (see Supplementary Figure. S7). Finally, the above procedure is combined with IG using various baselines and nonlinear paths between samples as we describe below.

**Figure 1.**
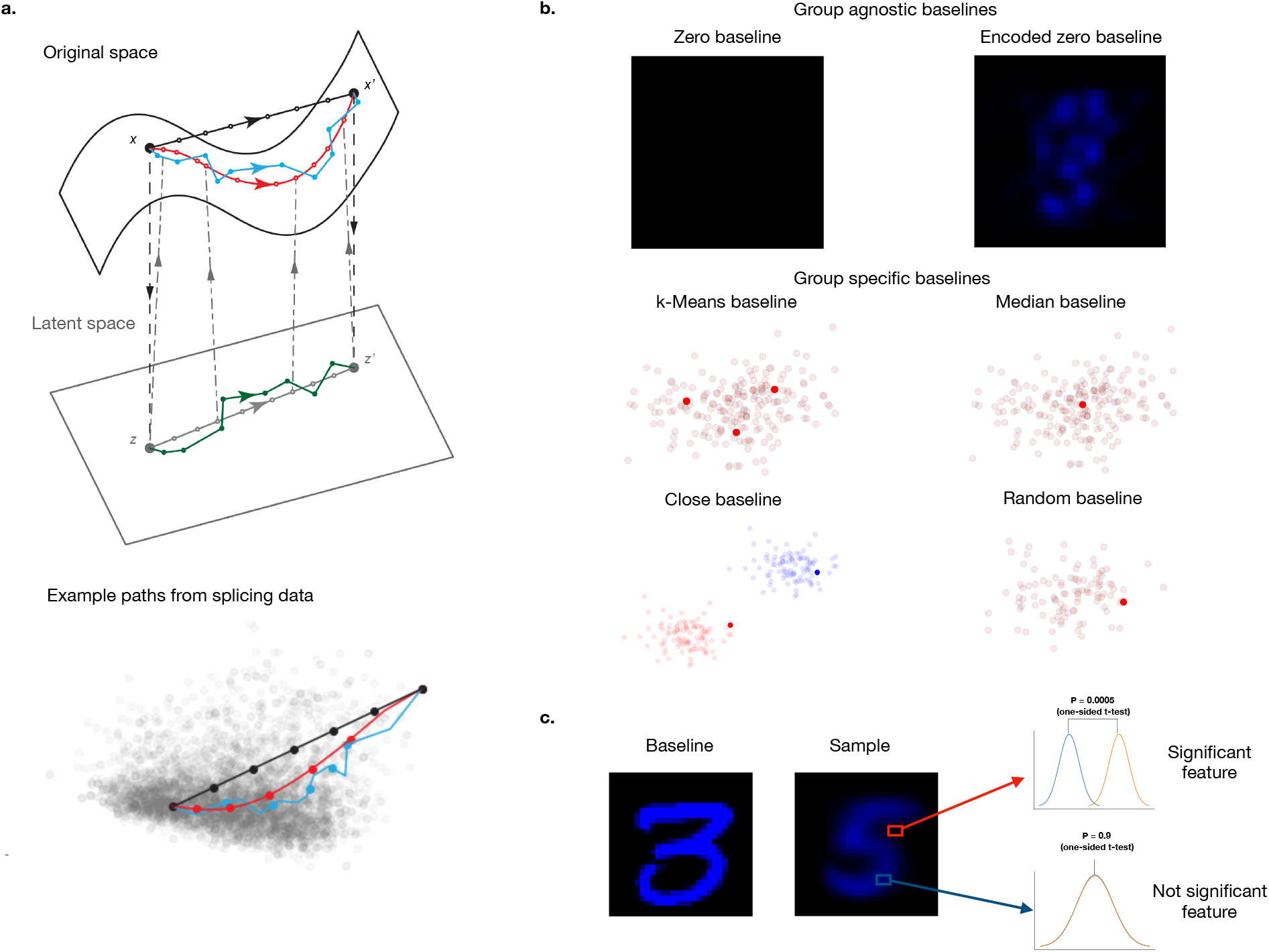
IG Enhancements to identify relevant features per sample or task. **a**, Top: Illustration of different linear and nonlinear paths for IG in the original. (1) Linear path in original feature space (**O-L-IG**, black line), (2) Neighbors path in the original feature space (**O-N-IG**, blue line), (3) Linear path in the hidden feature space (H-L-IG, gray line) and (4) Neighbors path in the hidden feature space (**H-N-IG**, green line). Bottom: Visualization of such paths for a specific sample using splicing data from (Keane et al., 2011) and PC1, PC2 of the feature space (see Supplementary Information Section 4). **b**, Illustration of different group agnostic (zero and encoded zero) and group specific (*k*-means, median, close, random) baselines. Encoded zero baseline is generated by decoding a zero vector in the latent space. A close baseline is created by taking a baseline point which is close to the sample in euclidean distance (see Methods). **c**, Illustration of framework to identify significant features that distinguish sample 5 from baseline 3. The digit images show mean of 300 examples of sample digit 5 and median of 300 examples of baseline digit 3. The distribution plots are illustrative only. They show difference between distribution of attributions of two sets of samples for a significant and a not significant feature.

### IG with nonlinear paths identify significant features missed using linear paths or simple gradients

Using the framework described above we first evaluated the effect of different nonlinear paths. While the original IG only considered linear paths in the original feature space, Fig. 1a shows several different paths evaluated in this work: (1) Linear path in original feature space (**O-L-IG**, black line), (2) *k*-nearest neighbors path in the original feature space (**O-N-IG**, blue line), (3) Linear path in the hidden feature space (**H-L-IG**, gray line) and (4) *k*-nearest neighbors path in the hidden feature space (**H-N-IG**, green line). Fig. 2a shows that simple gradients and the original IG as proposed in (Sundararajan et al., 2017), O-L-IG, fails to identify any known regulatory features as significant. In contrast, using nonlinear paths in O-N-IG, H-L-IG and H-N-IG, identify 488, 85, and 24 meta-features respectively.

**Figure 2.**
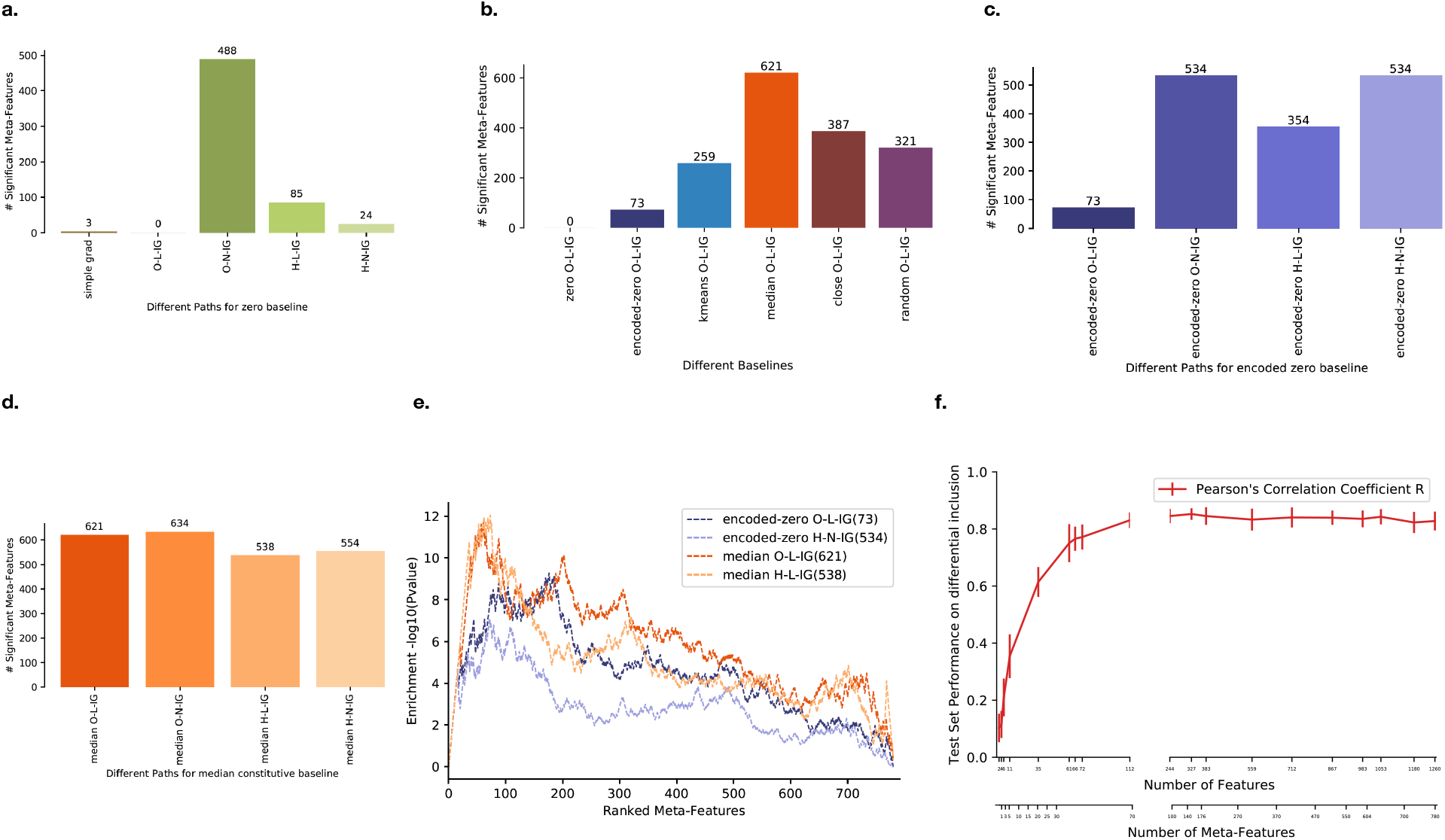
Performance evaluation of original IG and various enhancements on splicing data. **a**, Number of significant meta-features identified by different simple gradients, original IG (O-L-IG) and nonlinear paths (see main text) with zero baseline. **b**, Number of significant meta-features identified by different baselines (explained in text) with a linear path in the original feature space. **c**, Number of significant meta-features identified by different paths with an encoded-zero baseline. **d**, Number of significant meta-features identified by different paths with three median-constitutive baseline points. **e**, Enrichment of known brain regulatory features in significant features identified by the best two paths each from encoded-zero and median-constitutive baselines. **f**, Splicing prediction with increasing subsets of significant features identified by IG on latent-linear path with median baseline from constitutive splicing events. The x-axis shows both number of features and number of meta-features. Significant features passing one-sided t-test, Bonferroni adjusted *p*-value ≤ 0.05.

While the number of significant features identified by the various methods vary drastically, it is admittedly hard to assess levels of false positive features for such a real life prediction task or argue that identifying more features is strictly better. Nonetheless, the lack of significant features when using simple gradients or vanilla IG is striking. Furthermore, previous modeling efforts reported hundreds of those features as relevant and qualitatively, these features include many known regulatory features we expect to find such as conservation score and RNA binding proteins. The relevance of the identified features is also supported by known features enrichment analysis (see below).

### Group agnostic and group specific baselines identify significant features missed by a zero baseline

Next, we evaluated several approaches to define a reference point, or groups of those, for the different linear and nonlinear paths. First, we considered a *group agnostic* reference which does not require any prior biological information to define it. Specifically, an encoded-zero baseline (encoded-zero O-L-IG, Fig. 1b top panel) is generated by decoding the zero vector in the latent space of our autoencoder. This approach identifies 73 significant meta-features, a marked improvement over the original zero O-L-IG which failed to identify brain specific features. When the encoded-zero baseline is combined with nonlinear paths, substantially more significant features are identified (354-534) as shown in Fig. 2c.

We also evaluated several approaches to define a *group specific* baseline using different methods for selecting reference points (*k*-means, median, close and random) as shown in Fig. 1b, bottom panel. These baselines are designed to find significant features that distinguish differential inclusion in brain from constitutive splicing events. This means that the identified features should make a specific splicing event both an alternative exon (*e.g*., weak splice sites) and differentially included in brain (*e.g*., NOVA motif clusters). As expected, this refined definition of the baseline results in more significant meta-features being identified compared to those found by the group agnostic baseline, with median O-L-IG identifying the highest number of significant meta-features (621, Fig. 2b). Notably, we found that median baseline performs well across all linear and nonlinear paths (Fig. 2d, see Supplementary Figure S5 to see performance of other baselines). Here too the relevance of the identified features was assessed using known features enrichment analysis (see below) and comparison to a random sample group. Specifically, using a similar sized random group of events produced well calibrated *p*-values with no feature passing a *p*—value of 0.01 after multiple hypothesis correction (see Supplementary Figure. S7).

When we tested for the overlap of the features reported as significant by the various path and reference point definitions we found extensive overlap, indicating robustness (Supplementary Figure S6). For example, models that used a group agnostic baseline of encoded zero only had 4/534 shared features which were not reported by the models using a group specific baseline. To get a quantitative assessment of the biological relevance of the reported features, we tested them for enrichment of known splicing features and brain-specific regulatory features that are different from constitutive splicing events. Fig. 2e shows that significant meta-features identified by constitutive baselines with linear (median O-L-IG) and nonlinear (median H-L-IG) paths have higher enrichment of these known regulatory features than the *group agnostic* baseline (encoded-zero O-L-IG and encoded-zero H-N-IG) thus emphasizing the biological relevance of improving model interpretation by selecting meaningful baselines.

### Identified significant features improve splicing prediction

An important question related to the identified features relevance regards their effect on prediction. To address this question, we evaluated the role of significant meta-features for predicting ΔΨ on the set of differentially spliced exons in brain. Notably, this is a simpler problem than the splicing code prediction task and thus the prediction improvement for the differentially included exons in the test set saturates around 100 meta-features (244 original features) as seen in Fig. 2f. This early saturation is to be expected since splicing code features have information relevant to additional tasks such as identifying alternative exons and distinguishing low-inclusion and high-inclusion exons that are not needed for this task.

### Features distinguishing two digits in MNIST are located in key areas and allow accurate prediction

Finally, we evaluated our framework on the task of prediction attribution for handwritten digit recognition using the MNIST dataset (LeCun, 1998). We follow similar procedure as the splicing code model to create the joint model from a variational autoencoder and a feed-forward network. Using this network, we generate attributions for the digit 5 from the baseline digit 3. Fig. 3a, left panel shows mean attributions across 300 examples of 5 from median baseline 3 using the linear path in the hidden space. We can see that to distinguish the digit 5 from a 3, median H-L-IG identifies that the pixels on the bottom left and the top right should be absent (red) while the top left should be present (green). When combined with the significant features selection procedure these results become more pronounced, while other weaker attributions are removed (Fig. 3a, right panel). More generally, to test whether our framework identifies significant features that can distinguish the sample class from the baseline class, we train networks using only significant pixels (8% to 20% of all pixels) as features on the task of distinguishing the sample digit 5 with each remaining digit serving as baseline in a different network. Fig. 3b shows that the accuracy of this model on the test set is almost as good (0.0% to 1.44% loss in accuracy) as a network trained with all 784 pixels as features.

**Figure 3.**
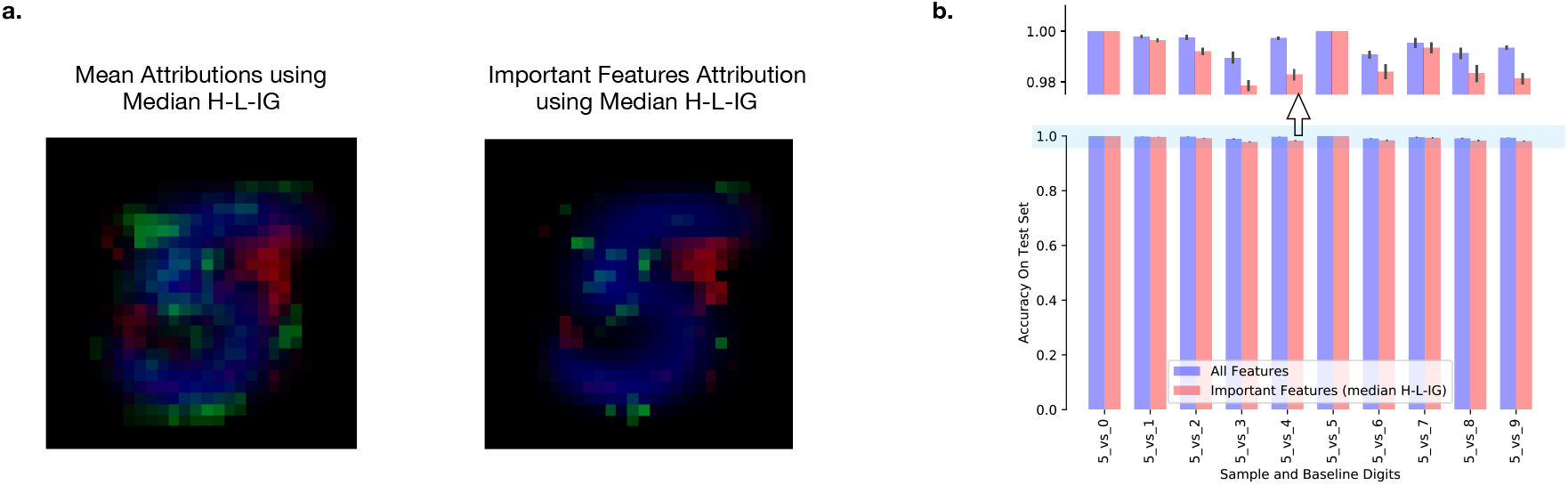
Extended IG framework with handwritten digit data. **a**, On the left, mean attributions generated from 300 examples of digit 5 and median baseline digit 3 using our approach. On the right, the subset of statistically significant features for the same set (one-sided t-test, Bonferroni adjusted *p*-value ≤ 0.05). Pixels belonging to the digit 5 are blue, positive attribution shown in green and negative attribution shown in red. **b**, Performance of models trained to distinguish sample from the baseline digits using all features and only the significant features identified using our approach (0.00% to 1.44% loss in accuracy while using 8% to 20% of all pixels). The top panel enlarges the y-axis (0.98 to 1.00) to highlight the differences in performance. These models solve the binary classification task of distinguishing the sample digit from the baseline digit and thus require fewer pixels than the original multi-class classification problem of classifying each image as 1 of 10 possible digits.

## Discussion

While the usage of deep learning in genomics has grown exponentially in the recent years (Ching et al., 2018), relatively little work has been done to assess or improve the interpretation of these models. We define model interpretation in terms of feature attribution *i.e*., finding features that are significant for a specific prediction task. We present a framework to identify such features, building upon the method of integrated gradients (IG) (Sundararajan et al., 2017). Our specific contributions are proposal of nonlinear paths and group agnostic and group specific baselines for integrated gradients resulting in improved feature detection, and the addition of a statistical test to assess significance of different features. Additionally, we ensure robustness of our results by assessing stability of data points in latent spaces of different model architectures via neighbor similarity.

Lack of gold standard is likely a major reason for the scarcity of published work on model interpretability in genomics. Simulated data or image recognition data, where the ground truth is readily available, is often used to get around this issue. For example, DeepLIFT, a notable effort for interpretation in genomics, uses simulated motif sequences and MNIST digit recognition data (Shrikumar et al., 2017). Here, we sidestep this issue by focusing on the splicing code prediction task which involves identifiable features. Admittedly, the evaluation of significant features for this real life task does not offer quantitative assessment of false positive or false negative features. In addition, our approach does not offer any theoretical guarantees beyond those of sensitivity and implementation invariance discussed before. Specifically, the usage of a reference sample has been shown to overcome issues of activation saturation (Shrikumar et al., 2017), but we lack theoretical guarantees for identifying significant features when they are highly correlated. Here, we compute the effect of highly correlated features by the maximal attribution assigned to a feature in that group (see Methods). More advanced approaches may be applied to handle this issue but in practice we found that even comparing the contribution of those redundant features to the null distribution allowed us identify them as individually significant, indicating robustness.

Despite the limitations discussed above, we were able to demonstrate that our approach controls for false positive features when compared to a randomly selected reference group, the identified features give high prediction accuracy when used alone, and that the enhancements we propose to IG enable it to identify many more biologically relevant features. Specifically, we find simple gradient and IG with zero baseline perform poorly. Instead, users in search of a group agnostic baseline can use an encoded-zero baseline. Of the nonlinear paths we offer, H-L-IG is the most robust across different baselines while being computationally more efficient than the O-N-IG and H-N-IG paths. However, connectivity in the latent space, as is observed in single cell data, may motivate neighbour based approaches in other applications. The performance of baseline selection methods for the group specific baseline depends on the underlying structure of the data but using the median points is computationally efficient and yields best results on the splicing data. Finally, we show our approach carries over to another more interpretable task in vision.

Overall, this work offers a framework for researchers in genomics to interpret deep learning models. We hope this framework, along with the available code, will help gain better biological insights from such models in the future.

## Methods

### Notation

We are interested in interpreting the prediction made by a deep learning model given an input observation by assigning attributions to each feature of the observation. Here, we assume that predictions are made from inputs ***x*** in a *p*-dimensional feature space 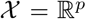. Predictions are obtained by a real-valued prediction function on feature space 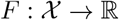. The goal is to be able to obtain a *p*-dimensional vector of attributions called **attr** ∈ ℝ^*p*^, with each coordinate corresponding to the *p* dimensions of the feature space, for an input ***x*** that indicates how each of the *p* features contributes to the prediction *F*(***x***).

### Datasets

In this paper, we use RNA-Seq experiments processed by (Keane et al., 2011) from six mouse tissues (Hippocampus, Heart, Liver, Lung, Spleen, Thymus) with average read coverage of 60 million reads. We generated 1,357 genomic features from 14,596 exon skipping events and 74,156 constitutive exon triplets using AVISPA (Barash et al., 2013). Ψ and ΔΨ quantification for the exon skipping events were generated using MAJIQ (Vaquero-Garcia et al., 2016). Furthermore, we use MNIST handwritten digits dataset (LeCun, 1998). This dataset contains 70,000 images of handwritten digits from 0 to 9. Each image contains 28 × 28 pixels.

### Splicing Code Model

We first train an autoencoder to find a latent embedding for the 1, 357 splicing features in a lower dimensional space. Subsequently, we use this embedding along with a separate input for two tissues types as the input to a feed-forward neural network. Predictions are made for three targets:

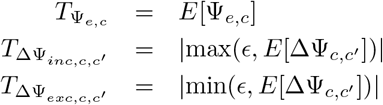

T_Ψ_*e,c*__ is the expected PSI value of the event *e* in condition *c*, T_ΔΨ_*inc,c,c*′__, captures the ΔΨ for events with increased inclusion between condition *c* and *c*′ and T_ΔΨ_*exc,c,c*′__ captures the ΔΨ for events with increased exclusion between condition *c* and *c*′. *ϵ* is a uniform random variable with values between 0.01 and 0.03. It is used to provide small ΔΨ values for non-changing events. Our goal is to interpret predictions made by this model, attributing which of the 1, 357 features contribute to *T*_Ψ_*e,c*__, *T*_ΔΨ_*inc,c,c*′__ or *T*_ΔΨ_*exc,c,c*′__ for a given sequence. Thus we combine the encoder from the autoencoder and the splicing code feed-forward network to form a single network. The attributions are computed on this combined network.

### Handwritten Digit Model

We use a feed-forward neural network architecture for this task with a variational autoencoder for dimensionality reduction. The input to the feed-forward neural network is the latent embedding from the variational autoencoder. Prediction is performed to identify the class of the input image from 10 digit classes. Interpretation task involves attributing predictions made by this model to a subset of 28 × 28 (784) pixels. The attributions are computed on the joint network combining the encoder from the variational autoencoder and the digits feed-forward network.

### Evaluation of Latent Space Stability

As is common in many genomic learning tasks, the splicing codes described above involves an embedding of the original features in a lower dimensional latent space. This embedding leads to an extension of IG by using this latent space, but at the same time raises the question of whether the embedding itself is robust. Lack of robustness in embeddings can lead to undesirable scenarios where different network architecture choices lead to different interpretations. One way to ensure robustness of different embeddings is to ensure similar relative distances between different data points in different embeddings. Thus, we compute the spearman rank correlation of the pairwise distances among training points in latent space between different autoencoders. We also evaluate correlations to the pairwise distances in the original feature space for comparison. Supplementary Figure S3 shows that different autoencoder and variational autoencoder embeddings are stable (spearman rank correlation between architectures ranges from 0.70 to 0.96).

### LIG: Latent Integrated Gradients

Integrated Gradients (Sundararajan et al., 2017) is a feature attribution method for explaining predictions from a differentiable function *F* such as is obtained from deep learning models. The per-feature attributions for a prediction are defined relative to a reference point 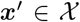 and its prediction *F*(***x***′). For an observation 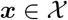, integrated gradients obtains an attribution vector **attr**(***x***) by integrating the gradient of *F* with respect to the feature space along a path 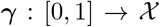 that starts at ***x***′ and ends at ***x***, i.e. *γ*(0) = ***x***′ and *γ*(1) = ***x***. If we write the attribution vector coordinate-wise as:

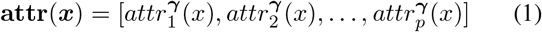

Then, the attribution for the *j*-th feature *x_j_* is:

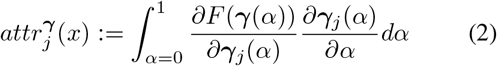

(Sundararajan et al., 2017) focuses on the special case where the path *γ* is chosen to take the straight-line path on ℝ^*p*^ from ***x***′ to ***x***. Parameterized by *α* ∈ [0,1], the path is *γ*(*α*) = ***x***′ + *α*(***x*** − ***x***′) so that the attribution for the *j*-th feature is:

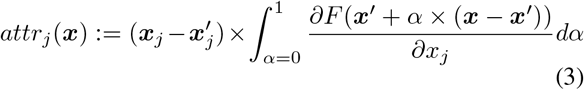

To address the possibility of artifactual attributions by integrated gradients from linear paths crossing regions of ℝ^*p*^ far from training points in feature space used to obtain *F*, we evaluate nonlinear paths designed to stay close to the data. We evaluate two approaches for generating such nonlinear paths: autoencoder networks on feature space and nearest neighbor graphs of training data. In the first approach, nonlinear paths are created by considering an autoencoder as the composition of an encoder 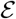 and decoder 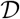 to and from some latent space 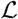. We define 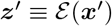 and 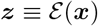. We take the linear path between ***z***′ and ***z*** and use 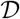 to map it back to a nonlinear path on 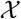. The final path accounts for the reconstruction error in mapping the endpoints back to 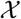 by connecting ***x***′ and ***x*** to the start and end of the decoded path (i.e. linearly interpolating between the endpoints and their auto-encoded counterparts).

In the second approach, nonlinear paths can be created in the original feature space 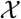 or the latent space 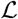. In 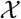, we construct the *k*-nearest neighbor graph on training data with respect to a distance metric on the 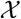, weighting edges by distances between points. The path between ***x***′ and ***x*** is created by adding them to the nearest-neighbors graph and finding the shortest path between them using Dijkstra’s algorithm, interpolating linearly between neighbors. In 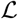, the procedure is similar except two key differences. First, the distances are computed on the 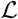. Second, the *k*-nearest neighbors path is computed between ***z***′ and ***z*** and the decoder 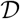 is used to map this path back to 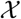. Computing the *k*-nearest neighbors graph can be computationally expensive making the linear paths in original and latent spaces computationally more efficient.

### Evaluation of IG based methods for feature attribution

The original IG work by (Sundararajan et al., 2017) only offered a method to compute the attribution per feature but did not offer a measure to identify significant features associated with a specific task. To address this need, we propose Algorithm 1. The input *G* contains attributions for a set of events of interest (e.g. differentially included exons in brain) and *R* contains attributions for a random set of events. For each feature, we perform a one-sided t-test for the positive tail of the distribution on *Set_G_* and *Set_R_*. The one-sided t-test captures features where absolute attribution on *Set_G_* is higher than *Set_R_*. To address the multiple-testing problem, we perform Bonferroni correction with family-wise error rate (FWER) of 0.01. A possible limitation of the above approach is that attribution may be dispersed between highly related features such as slight variations of the same splice factor binding motif. To address this issue in the context of splicing codes, we group highly-correlated features into 781 meta-features, as done in previous works (Barash et al., 2010; 2013). To compute the attribution for a meta-feature, we sum the attributions of all its features. Then we apply Algorithm 1 to find significant meta-features.

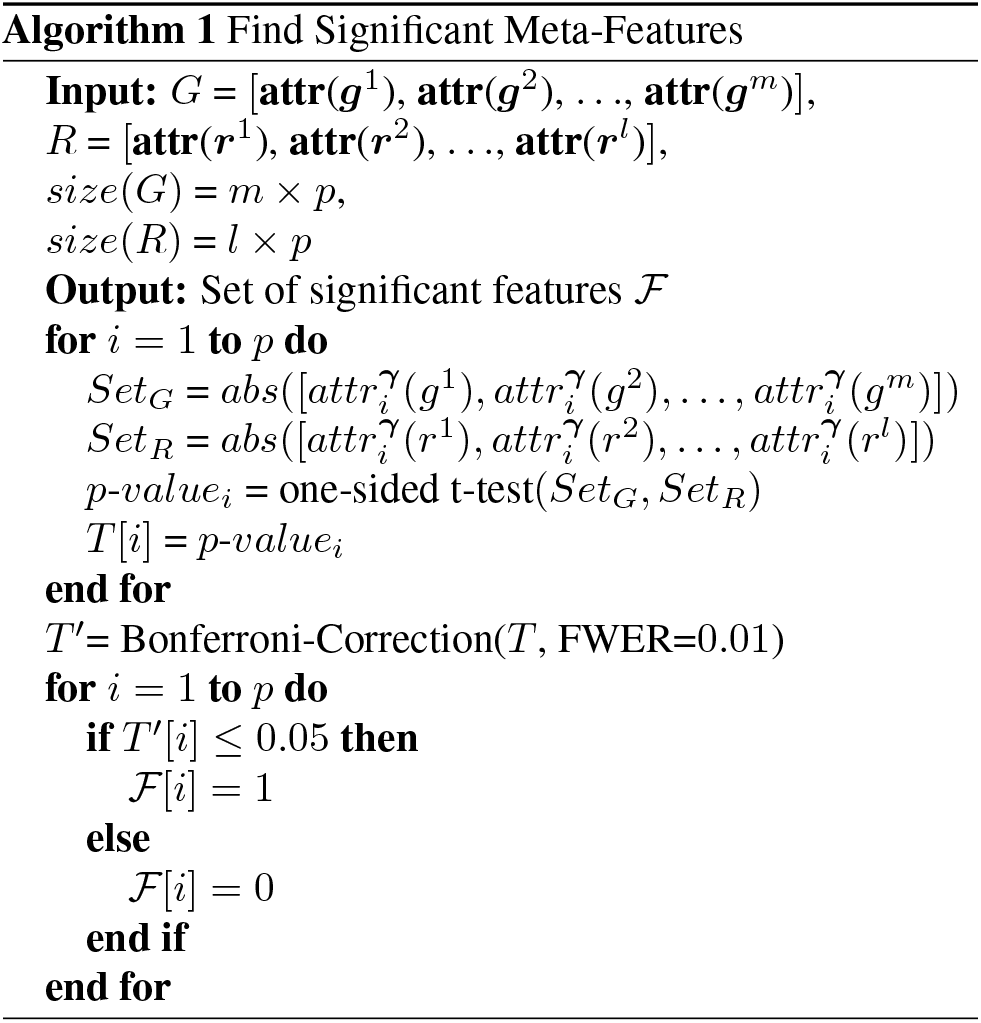

### Baselines

We use baselines to address two interesting questions about interpretation in genomics. An all-zero baseline is used to represent absence of signal in (Sundararajan et al., 2017). This is an acceptable choice for the object recognition task they work on since it represents an all black image. However, in the genomics domain, an all zero baseline is not meaningful. Therefore, the first question we address is: what is a good baseline to get meaningful interpretation for genomics? To answer this we propose a generic alternative baseline which we call **encoded-zero**. It requires an encoder/decoder to/from latent space such that we can use an all-zero point in the latent space and pass it to through the decoder to generate our baseline. The **encoded-zero** represents the mean of the data on which the autoencoder was trained. Interpretation with this baseline captures features that deviate from the mean and thus contribute to a sample’s prediction.

**Encoded-zero baseline** finds features that are important for a specific sample in comparison to the mean. However, in genomics, we are often interested in features that distinguish two classes of events. Thus, the second question we address is: how do we select baseline points from the reference class that can distinguish the class of interest from this reference class? We propose four approaches to select one or more baseline points. **Random baseline**: randomly sample one or more points from the baseline class. This serves as the naive method to evaluate the effectiveness of the other methods of selecting baselines. *k*-**means baseline**: cluster the points of the baseline class to *k* different clusters and then use cluster centroids as baseline points, the number of clusters can be selected by cross-validation, we observed that 3-5 clusters were enough for the splicing dataset. This method gives baseline points that represent different subgroups that might be present in the baseline class. **Median baseline**: compute euclidean distance of all the points of the baseline class from the median and select the points closest to the median. Points chosen using this method protect the later interpretation against outliers from the baseline class. **Close baseline**: compute euclidean distance of all the points in the baseline class from all the points in the class of interest and pick points from the baseline class that are close to a sample from the class of interest as its baseline. These baseline points are close to the sample and may represent minimal set of distinguishing features between the baselines and the points of interest. We discard the closest point from the baseline class to avoid extreme outlier points.

### Implementation Specifications

We implement these models and methods in Python using NumPy and Tensorflow. Paths are approximated discretely by a sequence of points on the path. Integration is performed numerically using the trapezoidal rule as implemented in NumPy. Approximation error of integration is estimated in two ways: (1) by comparing the sum of attributions to the difference between the predictions at the endpoints (as suggested by (Sundararajan et al., 2017)) and (2) by halving the step size for numerical integration to evaluate relative agreement between more refined approximations. Estimated integration error is used to determine the number of points necessary in the discrete paths (see Supplementary Figure S4).

## Acknowledgements

We would like to thank Barry Slaff for his insightful comments and feedback on an early draft of this manuscript. We also thank Caleb Radens and Matthew Gazzara for their constructive criticism of the manuscript.

## Availability of data and materials

http://majiq.biociphers.org/interpret-jha-el-al-2019/

## Author’s contributions

YB, AJ, and JKA developed the methods. DS developed the variational auto-encoder and helped with preliminary analysis. JKA developed the nearest neighbors path, the model stability tests, and the initial analysis of MNIST. AJ developed the code and performed all other experiments. AJ and YB wrote the manuscript with input from JKA. All authors read and approved the final manuscript.

