## Supplementary Materials for "Improving interpretability of deep learning models: splicing codes as a case study"

### SUPPLEMENTARY INFORMATION: IMPROVING INTERPRETABILITY OF DEEP LEARNING MODELS

**Anupama Jha[1], Joseph K. Aicher[2], Deependra Singh[1] & Yoseph Barash[1,2]**
[1]Department of Computer and Information Science, School of Engineering
[2]Department of Genetics, Perelman School of Medicine
University of Pennsylvania
Philadelphia, PA 19104, USA
{yosephb}@upenn.edu

#### 1 SPLICING CODE MODEL ARCHITECTURE

Supplementary Table S1: Autoencoder(AE) and Variational Autoencoder(VAE) Architecture

| Name | Hidden units and layers | Activation Function |
|---|---|---|
| AE200 | 1357, 850, 500, 200, 500, 850, 1357 | tanh |
| AE400 | 1357, 850, 500, 400, 500, 850, 1357 | tanh |
| AE500 | 1357, 850, 500, 850, 1357 | tanh |
| AE600 | 1357, 850, 600, 850, 1357 | tanh |
| VAE100 | 1357, 850, 500, 100, 500, 850, 1357 | tanh |
| VAE200 | 1357, 850, 500, 200, 500, 850, 1357 | tanh |
| VAE500 | 1357, 850, 500, 500, 500, 850, 1357 | tanh |

Supplementary Table S2: Feed Forward Network Architecture

| Name | Hidden units and layers | Activation Function |
|---|---|---|
| AE500-encoded Network | 500, 500, 200, 50, 6 | ReLU |

Supplementary Table S3: Combined Encoder Feed Forward Network Architecture

| Name | Hidden units and layers | Activation Function |
|---|---|---|
| Combined AE500-encoded Network | 1357, 850, 500, 500, 200, 50, 6 | ReLU |

#### 2 MNIST DIGIT MODEL ARCHITECTURE

Supplementary Table S4: Autoencoder(AE) and Variational Autoencoder(VAE) Architecture

| **Name** | **Hidden units and layers** | **Activation Function** |
|---|---|---|
| VAE50 | 784, 500, 500, 50, 500, 500, 784 | ELU, tanh |

Supplementary Table S5: Feed Forward Network Architecture

| **Name** | **Hidden units and layers** | **Activation Function** |
|---|---|---|
| VAE50-encoded Network | 50, 400, 200, 50, 10 | ReLU |

Supplementary Table S6: Combined Encoder Feed Forward Network Architecture

| **Name** | **Hidden units and layers** | **Activation Function** |
|---|---|---|
| Combined VAE50-encoded Network | 784, 500, 500, 50, 400, 200, 50, 10 | ELU, tanh, ReLU |

#### 3 PATH INTERPOLATION

The number of points used to interpolate paths between a sample and a baseline was selected by estimating the relative numerical integration error for paths between randomly sampled pairs of points. **Relative error** was estimated in two ways: (1) **sum error**, and (2) **median reinterpolation error**.

Sum error is described in (1) and uses the property:

$$\begin{aligned}\sum \text{attributions}(f, \gamma[a \to b]) &= \sum \int_{\gamma[a \to b]} \nabla f \, \mathrm{d}\gamma \\ &= \int_{\gamma[a \to b]} \sum \nabla f \, \mathrm{d}\gamma \\ &= \int_{\gamma[a \to b]} \nabla f \cdot \mathrm{d}\gamma \\ &= f(b) - f(a)\end{aligned}$$

We define the sum error as the relative difference between the left-hand-side and the right-hand-side of the equation, using the numerically calculated attributions for the left hand side. Specifically:

$$\text{sum error (relative)} = \frac{|\text{RHS} - \text{LHS (numerical)}|}{\varepsilon + |\text{RHS}|}.$$

Median reinterpolation error is obtained by comparing attribution estimates from a path with the given number of points to a refined path with additional points interpolating between the original points. For a given feature, we say that $g^*$ is the true attribution which is being estimated numerically by $g(n)$ for a path with $n$ points. Then, reinterpolating with $r$ points, we estimate the relative error for the feature attribution as:

$$2 \times \frac{|g(n) - g(rn)|}{\varepsilon + \max(|g(n)|, |g(rn)|)}.$$

This gives a relative error estimate per feature for a path between points. The median reinterpolation error is the median of these estimates across features.

We calculated these estimates for 200 pairs of randomly selected points in our dataset using linear paths, comparing 50, 100, 250, 500, and 1000 points in the path. We found that 51% of the replicates have zero relative error regardless of estimation method or number of points, and 2% of the replicates have outlying sum errors greater than $10^5$ due to a negligible difference between predictions for the source and destination points. Plots of relative error (sum error and median interpolation error) are found in Supplementary Figure S4. We decided to use 250 points for our subsequent experiments as it had an acceptable relative error.

#### 4 Plotting details for real paths on splicing data (Fig 1a, bottom panel )

The plot shows PC1 (horizontal axis) versus PC2 (vertical axis) of original feature space($1,357$ features) trained on splicing data. The scattered gray points are subset of input data in the PC space. Black, red, blue paths show linear, latent-linear, and neighbors paths between the same source and destination points (matching conceptual/illustrative figure 1a, top panel). Source point is picked randomly and the destination point is picked to maximize distance from source point in PC space (10 components). Neighbors path is approximation, i.e., computing neighbors distances on first 10 principal components due to computational overhead. Random seeds were manually selected to highlight differences between linear and latent linear paths.

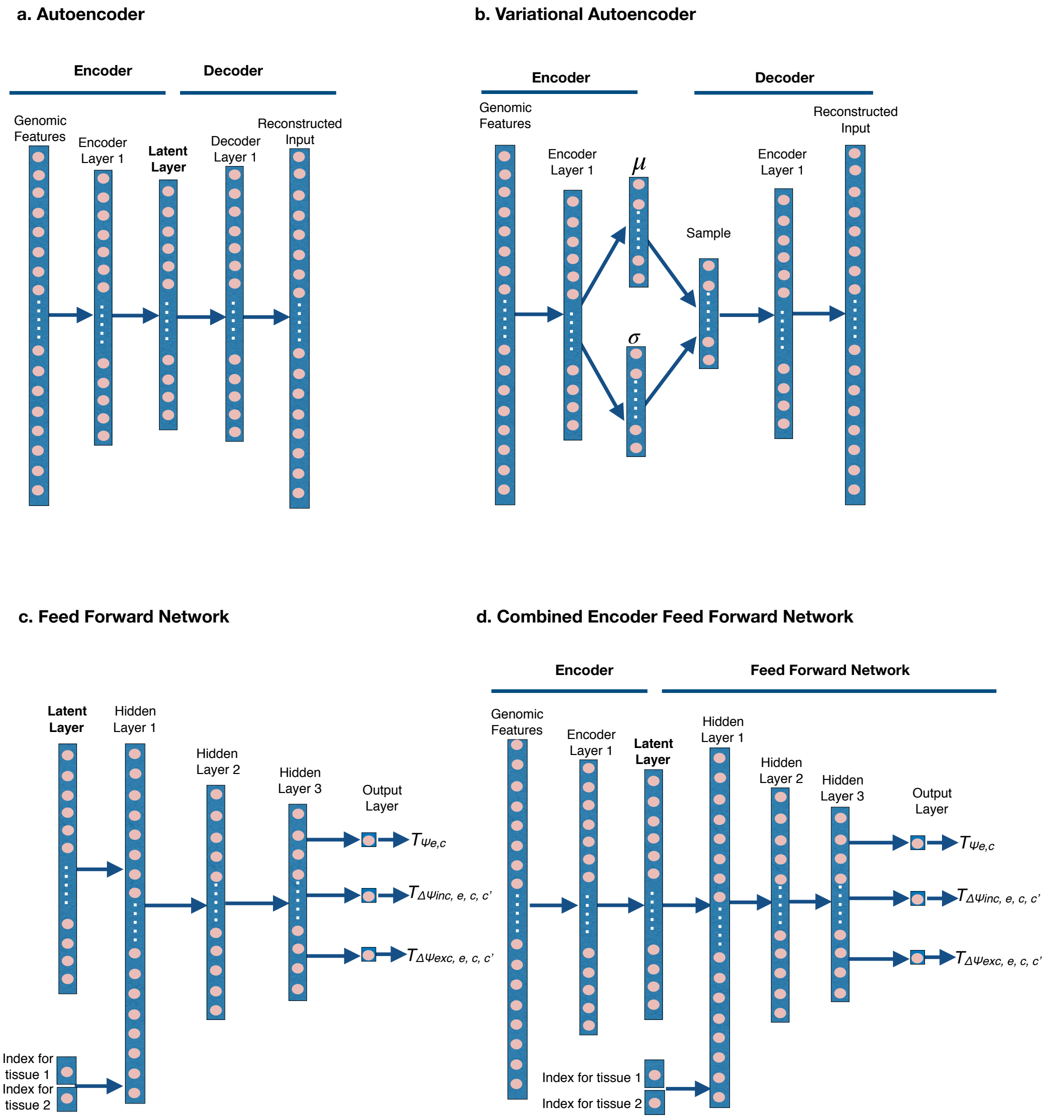


Supplementary Figure S1: **Splicing code architecture.** **a,** Illustration of the autoencoder architecture for the splicing code. **b,** Illustration of the variational autoencoder architecture for the splicing code. The latent layer learns mean $\mu$ and standard deviation $\sigma$ of the a Gaussian distribution from which samples are drawn for the decoder. **c,** Illustration of the Feed forward network for the splicing code. The latent layer from **a** is input to the model. The output of the model contains three targets: $\mathrm{T}_{\Psi_{e,c}}$ is the expected PSI value of the event $e$ in condition $c$, $\mathrm{T}_{\Delta\Psi_{inc,c,c'}}$ captures the dPSI for events with increased inclusion between condition $c$ and $c'$ and $\mathrm{T}_{\Delta\Psi_{exc,c,c'}}$ captures the dPSI for events with increased exclusion between condition $c$ and $c'$. **d,** Combined network from the encoder of the autoencoder in **a** and the feed forward network from **c**.

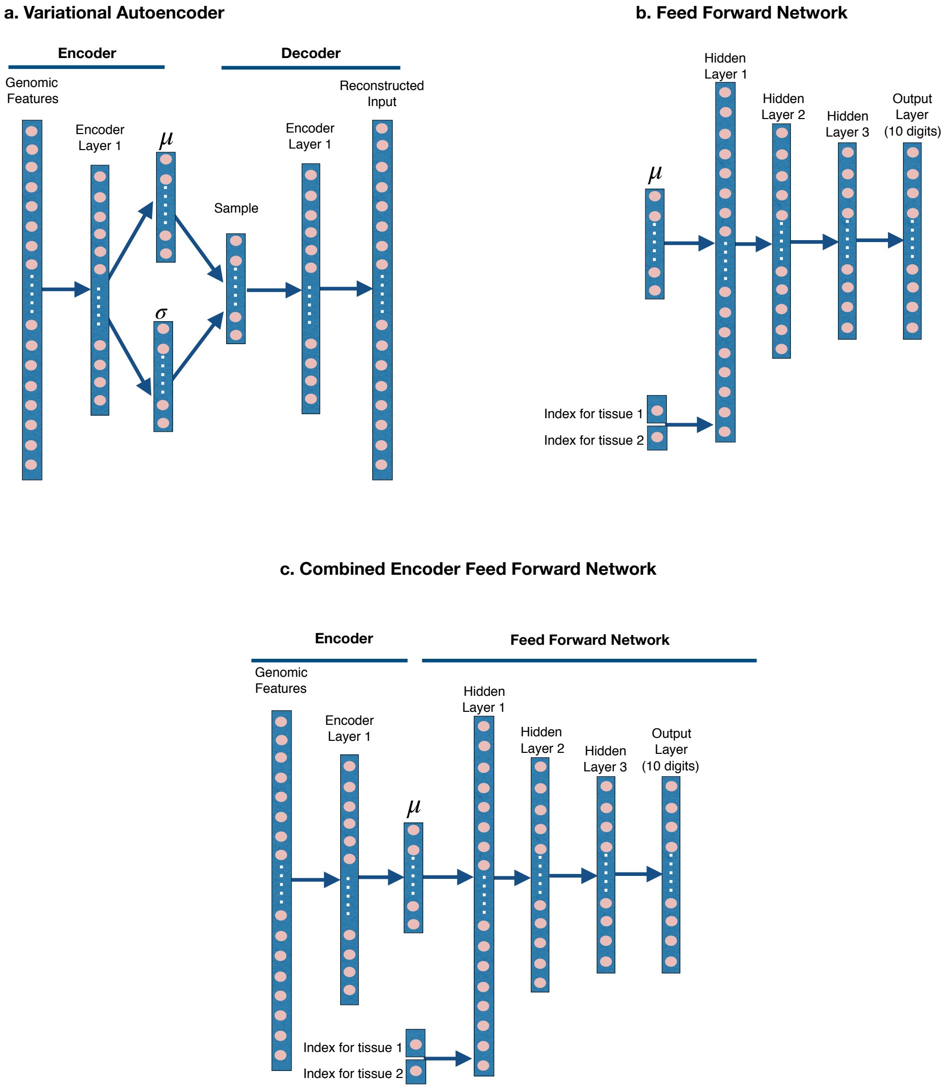


Supplementary Figure S2: **MNIST handwritten digit architecture. a,** Illustration of the variational autoencoder architecture for the digits model. The latent layer learns mean $\mu$ and standard deviation $\sigma$ of the a Gaussian distribution from which samples are drawn for the decoder. **b,** Illustration of the Feed forward network for MNIST handwritten digit task. The mean layer $\mu$ from **a** is input to the model. The output is a softmax over 10 digits. **c,** Combined network from the encoder of the variational autoencoder in **a** and the feed forward network from **b**.

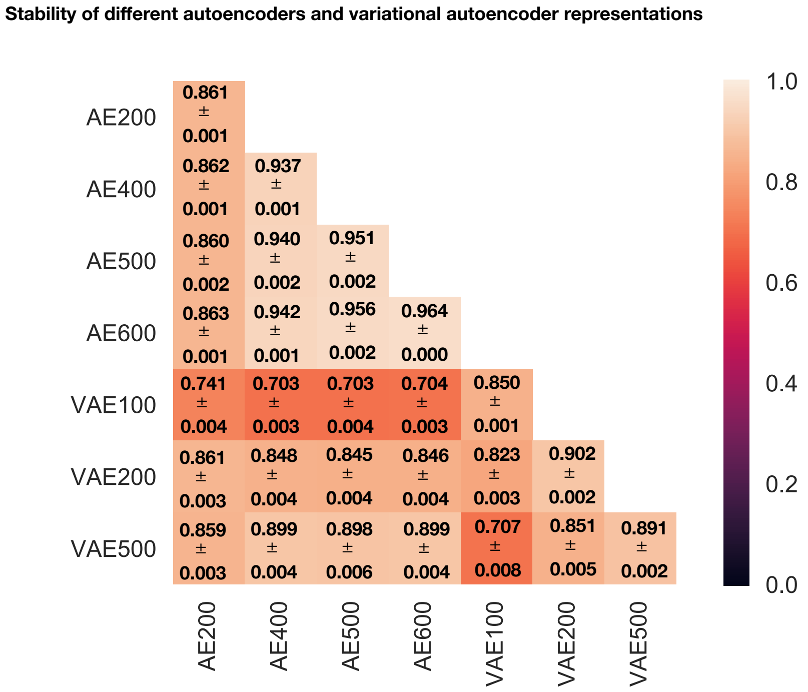


Supplementary Figure S3: **Stability analysis of autoencoders and variational autoencoders for the splicing code.** Spearman rank correlation of the pairwise distances among training points in latent space between different autoencoders and variational autoencoders. The distances are calculated for three randomly selected subset of training points to calculate the standard error. Supplementary Table S1-S6 describe the architecture of these networks.

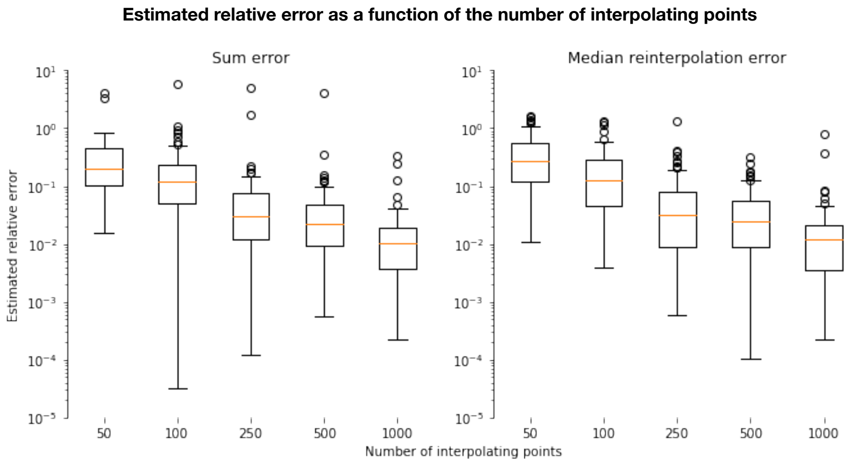


Supplementary Figure S4: **Estimated relative error as a function of the number of interpolating points**. The plots show relative error, as estimated by **(left)** sum error and **(right)** median reinterpolation error, versus an increasing number of points used to create a path between points. The relative error distributions are estimated over 200 pairs of randomly sampled points in the data, interpolated by each of the described number of points. 51% of the replicates have zero relative error regardless of estimation method or number of points and are excluded from the plot. 2% of the replicates have outlying sum errors greater than $10^5$ due to a negligible difference between predictions for the source and destination points and are excluded from the plot.

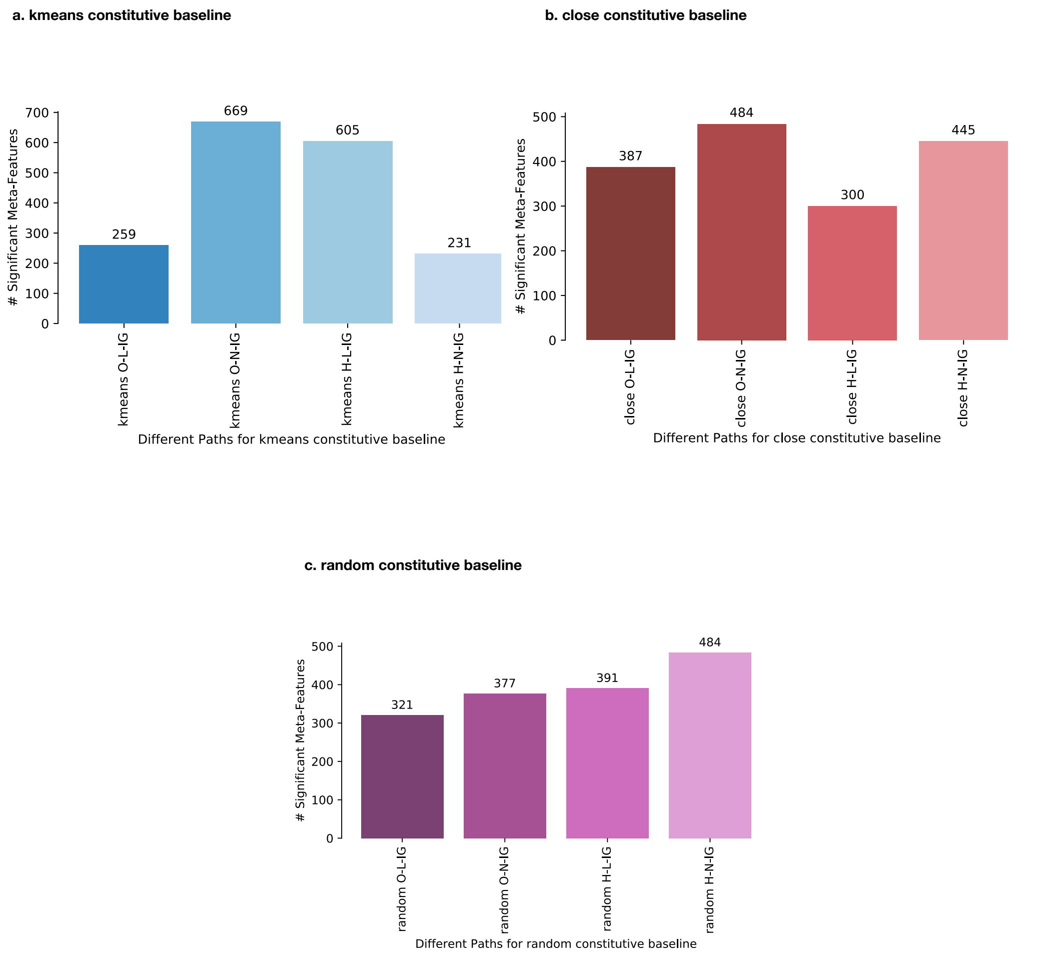


Supplementary Figure S5: **The effect of different paths and baselines on the number of significant features identified.** **a**, Number of significant meta-features identified by different paths with three kmeans-constitutive baseline points. **b**, Number of significant meta-features identified by different paths with three closest-constitutive baseline points. **c**, Number of significant meta-features identified by different paths with three random-constitutive baseline points.

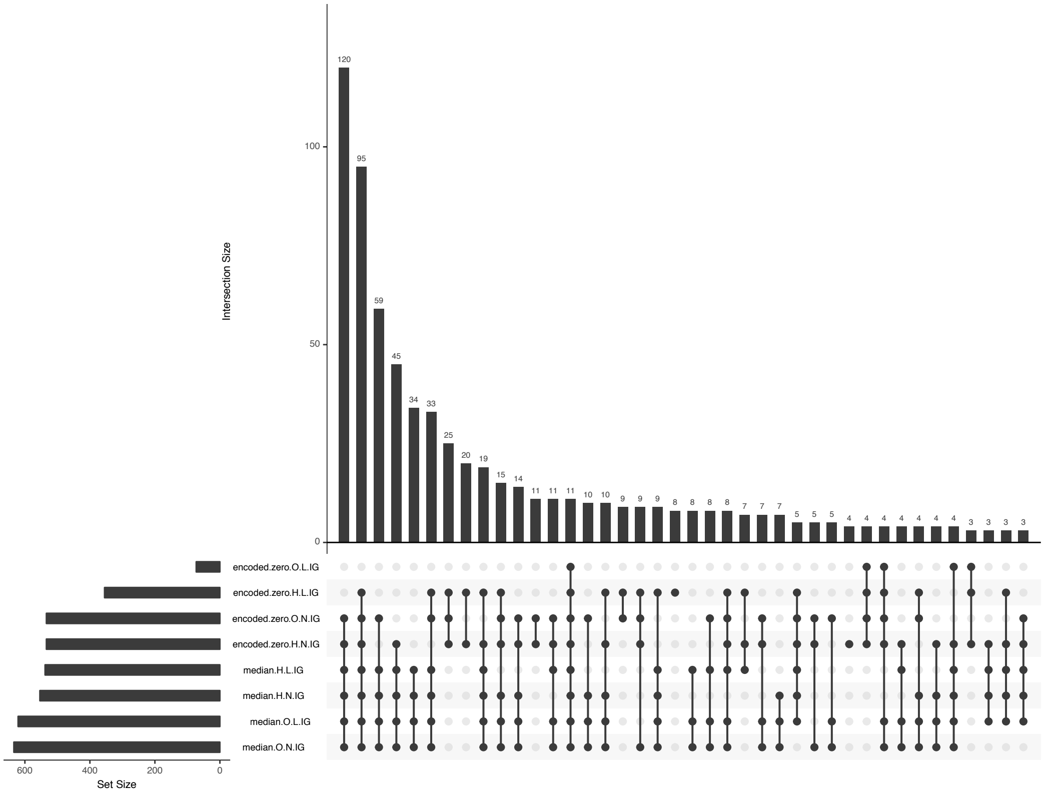


Supplementary Figure S6: **Overlap of significant features found by encoded-zero and median baselines.** The plot shows various intersections of features identified as significant using O-L-IG, O-N-IG, H-L-IG and H-N-IG paths for encoded-zero and median-constitutive baselines.

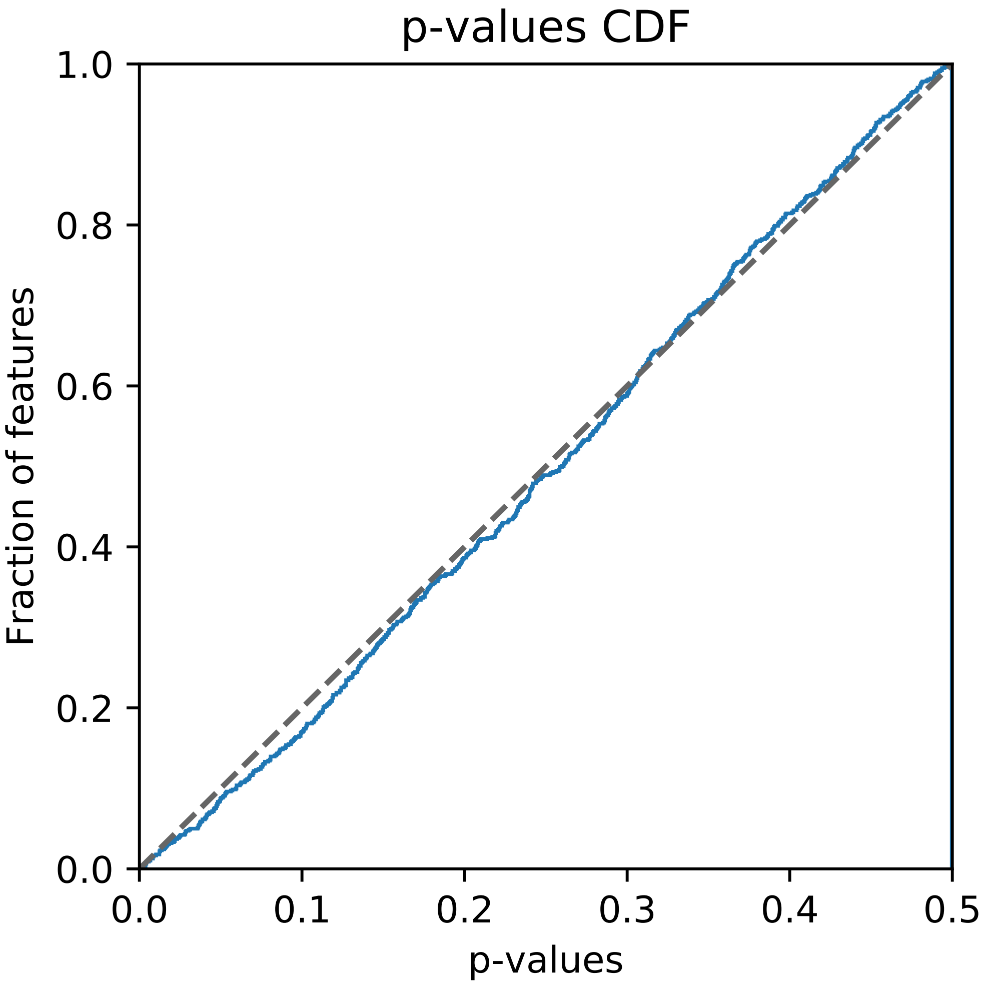


Supplementary Figure S7: **Calibration of *p*-value distribution**. Cumulative distribution of $p$-values for all meta-features using two random group of splicing events (one-sided t-test; $n = 781$ pairwise comparisons). The distribution of p-values is close to random (diagonal), indicating that empirical distribution of $p$-values is well calibrated.
